# WASABI: a dynamic iterative framework for gene regulatory network inference

**DOI:** 10.1101/292128

**Authors:** Arnaud Bonnaffoux, Ulysse Herbach, Angélique Richard, Anissa Guillemin, Sandrine Giraud, Pierre-Alexis Gros, Olivier Gandrillon

## Abstract

Inference of gene regulatory networks from gene expression data has been a long-standing and notoriously difficult task in systems biology. Recently, single-cell transcriptomic data have been massively used for gene regulatory network inference, with both successes and limitations. In the present work we propose an iterative algorithm called WASABI, dedicated to inferring a causal dynamical network from time-stamped single-cell data, which tackles some of the limitations associated with current approaches. We first introduce the concept of waves, which posits that the information provided by an external stimulus will affect genes one-by-one through a cascade, like waves spreading through a network. This concept allows us to infer the network one gene at a time, after genes have been ordered regarding their time of regulation. We then demonstrate the ability of WASABI to correctly infer small networks, which have been simulated *in silico* using a mechanistic model consisting of coupled piecewise-deterministic Markov processes for the proper description of gene expression at the single-cell level. We finally apply WASABI on *in vitro* generated data on an avian model of erythroid differentiation. The structure of the resulting gene regulatory network sheds a fascinating new light on the molecular mechanisms controlling this process. In particular, we find no evidence for hub genes and a much more distributed network structure than expected. Interestingly, we find that a majority of genes are under the direct control of the differentiation-inducing stimulus. In conclusion, WASABI is a versatile algorithm which should help biologists to fully exploit the power of time-stamped single-cell data.

## Introduction

It is widely accepted that the process of cell decision making results from the behavior of an underlying dynamic gene regulatory network (GRN) [1]. The GRN maintains a stable state but can also respond to external perturbations to rearrange the gene expression pattern in a new relevant stable state, such as during a differentiation process. Its identification has raised great expectations for practical applications in network medicine [2] like somatic cells [3–5] or cancer cells reprogramming [6,7]. The inference of such GRNs has, however, been a long-standing and notoriously difficult task in systems biology.

GRN inference was first based upon bulk data [8] using transcriptomics acquired through micro array or RNA sequencing (RNAseq) on populations of cells. Different strategies has been used for network inference including dynamic Bayesian networks [9, 10], boolean networks [11–13] and ordinary differential equations (ODE) [14] which can be coupled to Bayesian networks [15].

More recently, single-cell transcriptomic data, especially RNAseq [16], have been massively used for GRN inference (see [17,18] for recent reviews). The arrival of those single-cell techniques led to question the fundamental limitations in the use of bulk data. Observations at the single-cell level demonstrated that any and every cell population is very heterogeneous [19–21]. Two different interpretations of the reasons behind single-cell heterogeneity led to two different research directions:

1. In the first view, this heterogeneity is nothing but a noise that blurs a fundamentally deterministic smooth process. This noise can have different origins, like technical noise (“dropouts”) or temporal desynchronization as during a differentiation process. This view led to the re-use of the previous strategies and was at the basis of the reconstruction of a “pseudo-time” trajectory (reviewed in [22]). For example, SingleCellNet [23] and BoolTraineR [24] are based on boolean networks with preprocessing for cell clustering or pseudo-time reconstruction. Such asynchronous Boolean network models have been successfully applied in [25]. Other probabilistic algorithms such as SCOUP [26], SCIMITAR [27] or AR1MA1-VBEM [28] also use pseudo-time reconstruction complemented with correlation analysis. ODE based methods can be exemplified with SCODE [29] and InferenceSnapshot [30] algorithms which also use pseudo-time reconstruction.

2. The other view is based upon a representation of cells as dynamical systems [31,32]. Within such a frame of mind, “noise” can be seen as the manifestation of the underlying molecular network itself. Therefore cell-to-cell variability is supposed to contain very valuable information regarding the gene expression process [33]. This view was advocated among others by [34], suggesting that heterogeneity is rooted into gene expression stochasticity, and that cell state dynamic is a highly stochastic process due to bursting that jumps discontinuously between micro-states. Dynamic algorithms like SINCERITIES [35] are based upon comparison of gene expression distributions, incorporating (although not explicitly) the bursty nature of gene expression. We have recently described a more explicit network formulation view based upon the coupling of probabilistic two-state models of gene expression [36]. We devised a statistical hidden Markov model with interpretable parameters, which was shown to correctly infer small two-gene networks [36].

Despite their contributions and successes, all existing GRN inference approaches are confronted to some limitations:

1. The inference of interactions through the calculation of correlation between gene expression, whether based upon or linear [27] or non-linear [26] assumptions, is problematic. Such correlations can only reproduce events that have been previously observed. As a consequence, predictions of GRN response to new stimulus or modifications is not possible. Furthermore, correlation should not be mistaken for causality. The absence of causal relationship severely hampers any predictive ability of the inferred GRN.

2. The very possibility of making predictions relies upon our ability to simulate the behavior of candidate networks. This implicitly implies that network topologies are explicitly defined. Nevertheless, several inference algorithms [27–29, 35] propose a set of possible interactions with independent confidence levels, generally represented by an interaction matrix. The number of possible actionable networks deduced from combining such interactions is often too large to be simulated.

3. Regulatory proteins within a GRN are usually restricted to transcription factors (TF), like in [24, 26–30]. Possible indirect interactions are completely ignored. A trivial example is a gene encoding a protein that induces the nuclear translocation of a constitutive TF. In this case, the regulator gene will indirectly regulate TF target genes, and its effect will be crucial in understanding the GRN behavior.

4. Most single-cell inference algorithms rely upon the use of a single type of data, namely transcriptomics. By doing so, they implicitly assume protein levels to be positively correlated with RNA amounts, which has been proven to be wrong in case of post-translational regulation (see [33] for an illustration in circadian clock). Besides, at single-cell scale, mRNA and proteins typically have a poor linear correlation [34], even in the absence of post-translational regulation.

5. The choices of biological assumptions are also important for the biological relevance of GRN models. The use of statistical tools can be really powerful to handle large-scale network inference problem with thousand of genes, but the price to pay is loss of biological representativeness. By definition a model is a simplification of the system, but when simplifying assumptions are induced by mathematical tools, like linear [27–29, 35] or binary (boolean) requirements [23, 24], the model becomes solvable at the expense of its biological relevance.

In the present work we address the above limitations and we propose an iterative algorithm called WASABI, dedicated to inferring a causal dynamical network from time-stamped single-cell transcriptomic data, with the capability to integrate protein measurements. In the first part we present the WASABI framework which is based upon a mechanistic model for gene-gene interactions [36]. In the second part we benchmark our algorithm using *in silico* GRNs with realistic gene parameter values. Finally we apply WASABI on our *in vitro* data [37] and analyze the resulting GRN candidates.

## Results

Our goal is to infer causalities involved in GRN through analysis of dynamic multi-scale/level data with the help of a mechanistic model [36]. We first present an overview of the WASABI principles and framework. We then benchmark its ability to correctly infer *in silico*-generated toy GRNs. Finally, we apply WASABI on our *in vitro* data on avian erythroid differentiation model [38] to generate biologically relevant GRN candidates.

### WASABI inference principles and implementation

WASABI is a framework built on a novel inference strategy based on the concept of “waves”. We posit that the information provided by an external stimulus will affect genes one-by-one through a cascade, like waves spreading through a network (Fig 1-A). This wave process harbors an inertia determined by mRNA and protein half-lives which are given by their degradation rate.

By definition, causality is the link between cause and consequence, and causes always precede consequences. This temporal property is therefore of paramount importance for causality inference using dynamic data. In our mechanistic and stochastic model of GRN [36] (detailed in Method section Fig 7), the cause corresponds either to the protein of the regulating gene or a stimulus, which level modulates as a consequence the promoter state switching rates *k*_on_ (i.e. probability to switch from inactive to active state) and *k*_off_ (active to inactive) of the target gene. A direct consequence of causality principle for GRNs is that a dynamical change in promoter activity can only be due to a previous perturbation of a regulating protein or stimulus. For example, assuming that the system starts at a steady-state, early activated genes (referred to as early genes) can only be regulated by the stimulus, because it is the only possible cause for their initial evolution. An illustration is given in Fig 1-A: gene *A* initial variation can only be due to the stimulus and not by the feedback from gene *C*, which will occur later. A generalization of these concepts is that for a given time after the stimulus, we can infer the subnetwork composed exclusively by genes affected by the spreading of information up to this time. Therefore we can infer iteratively the network by adding one gene at a time (Fig 1-D) regarding their promoter wave time order (Fig 1-B) and comparing with protein wave time of previous added genes (Fig 1-C).

For this, we need to estimate promoter and protein wave times for each gene and then sort them by promoter wave time. We define the promoter activity level by the *k*_on_/(*k*_on_ + *k*_off_) ratio, which corresponds to the local mean active duration (Fig 1-B). Promoter wave time is defined as the inflection time point of promoter activity level where 50% of evolution between minimum and maximum is reached. Since promoter activity is not observable, we estimate the inflection time point of mean RNA level from single-cell transcriptomic kinetic data [37], and retrieve the delay induced by RNA degradation to deduce promoter wave time. Protein wave times correspond to the inflection point of mean protein level, which can be directly observed with our proteomic data [39]. A detailed description of promoter and protein wave time estimation can be found in the Method section. One should note that a gene can have more than one wave time in case of non monotonous variation of promoter activity, due to feedbacks (like gene A in our example) or incoherent feed-forward loop.

The WASABI inference process (Fig 1-C) takes advantage of the gene wave time sorting by adopting a divide and conquer strategy. We remind that a main assumption of our interaction model is the separation between mRNA and protein timescales [36]. As a consequence, for a given interaction between a regulator gene and a regulated gene, the regulated promoter wave time should be compatible with the regulator protein wave time. At each step, WASABI proposes a list of possible regulators in order to reduce the dimension of the inference problem. This list is limited to regulators with compatible protein wave time within the range of 30 hours before and 20 hours after the promoter wave time of the added regulated gene. This constraint has been set up from *in silico* study (see next section). For example, in Fig 1, gene *B* can be regulated by gene *A* or *D* since their protein wave time are close to gene *B* promoter wave time. Gene *C* can be regulated by gene *B* or *D*, but not *A* because its protein wave time is too earlier compared to gene *C* promoter wave time.

**Fig 1.**
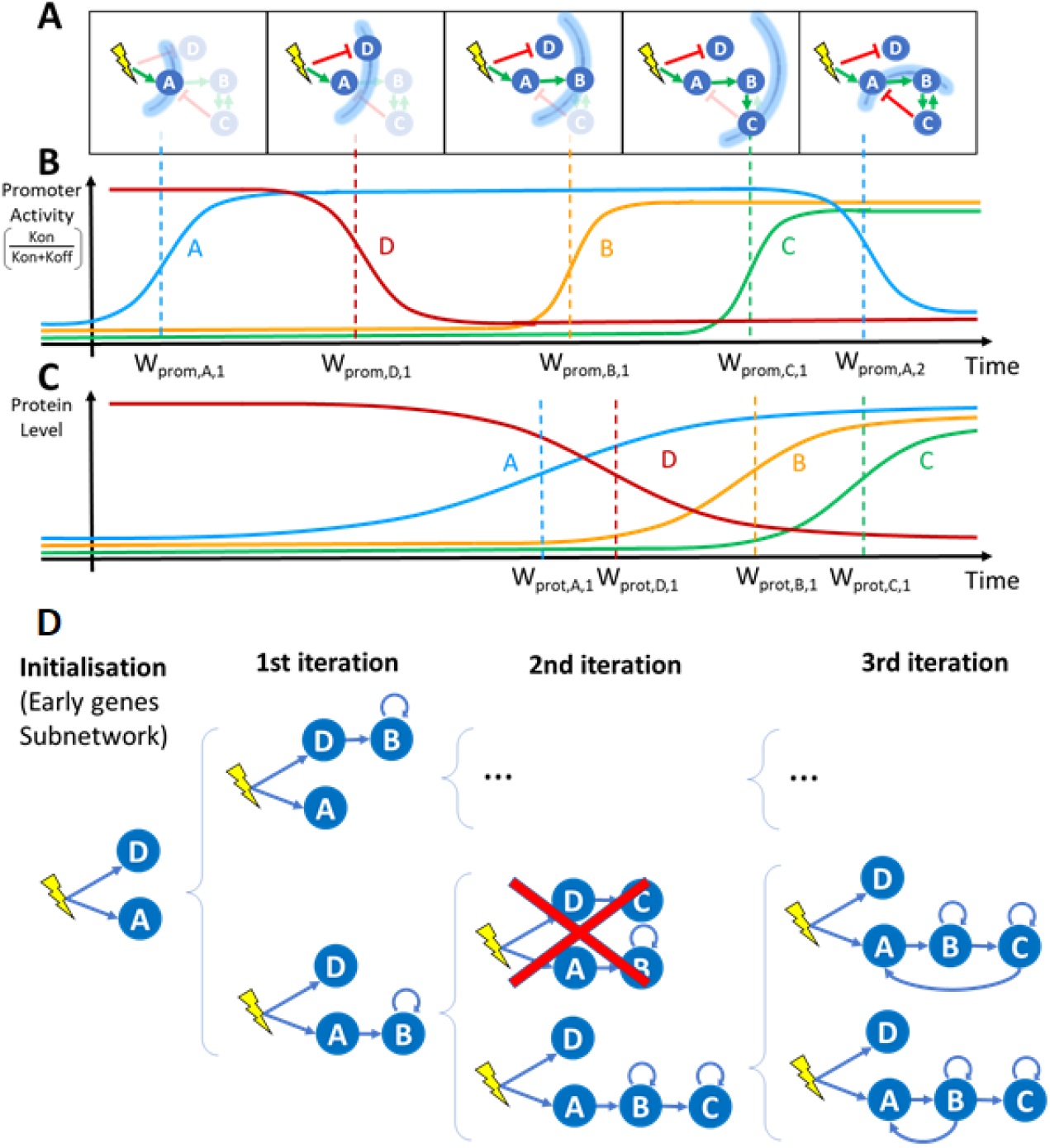
WASABI at a glance. A) Schematic view of a GRN: the stimulus is represented by a yellow flash, genes by blue circles and interactions by green (activation) or red (inhibition) arrows. The stimulus-induced information propagation is represented by blue arcs corresponding to wave times. Genes and interactions that are not affected by information at a given wave time are shaded. At wave time 5, gene *C* returns information on gene *A* and *B* by feedback interaction creating a backflow wave. B) Promoter wave times: Promoter wave times correspond to inflections point of gene promoter activity defined as the *k*_on_/(*k*_on_ + *k*_off_) ratio. C) Protein wave times: Protein wave times correspond to inflections point of mean protein level. D) Inference process. Blue arrows represent interactions selected for calibration. Based on promoter waves classification genes are iteratively added to sub-GRN previously inferred to get new expanded GRN. Calibration is performed by comparison of marginal RNA distributions between *in silico* and *in vitro* data. Inference is initialized with calibration of early genes interaction with stimulus, which gives initial sub-GRN. Latter genes are added one by one to a subset of potential regulators for which a protein wave time is close enough to the added gene promoter wave time. Each resulting sub-GRN is selected regarding its fit distance to *in vitro* data. If fit distance is too important sub-GRN can be eliminated (red cross). An important benefit of this process is the possibility to parallelize the sub-GRN calibrations over several cores, which results in a linear computational time regarding the number of genes. Note that only a fraction of all tested sub-GRN is shown.

For new proposed interactions, a typical calibration algorithm can be used to finely tune interaction parameter in order to fit simulated mRNA marginal distribution with experimental marginal distribution from transcriptomic single-cell data. To avoid over-fitting issues, only efficiency interaction parameter *θ*_*i,j*_ (Fig 7) is tuned. To estimate fitting quality we define a GRN fit distance based on the Kantorovitch distances between simulated and experimental mRNA marginal distributions (please refer to Method section for a detailed description of interaction function and calibration process). If the resulting fitting is judged unsatisfactory (i.e. GRN fit distance is greater than a threshold), the sub-GRN candidate is pruned. For genes presenting several waves, like gene *A*, each wave will be separately inferred. For example, gene *A* initial increase is fitted during initialization step, but only the first experimental time points during promoter activity increase will be used for calibration. Genes *B* and *C* regulated after gene *A* up-regulation will be added to expand sub-GRN candidates. Finally, the wave corresponding to gene *A* down-regulation is then fitted considering possible interactions with previously added genes (namely gene *B* and C), which permits the creation of feedback loops or incoherent feed-forward loops.

Positive feedback loops cannot be easily detected by wave analysis because they only accelerate, and eventually amplify, gene expression. Yet, their inference is important for the GRN behavior since they create a dynamic memory and, for example, may thus participate to irreversibility of the differentiation process. To this end, we developed an algorithm to detect the effect of positive feedback loops on gene distribution before the iterative inference (see Supporting information). We modeled the effect of positive feedback loops by adding auto-positive interactions. Note that such a loop does not necessarily mean that the protein directly activates its own promoter: it simply means that the gene is influenced by a positive feedback, which can be of different nature. For example, in the GRN presented in Fig 1-A, genes *B* and *C* mutually create a positive feedback loop. If this positive feedback loop is detected we consider that each gene has its own auto-positive interaction as illustrated in Fig 1-C. Positive feedback loops could also arise from the existence of self-reinforcing open chromatin states [40] or be due to the fact that binding of one TF can shape the DNA in a manner that it promotes the binding of the second TF [41].

### *In silico* benchmarking

We decided to first calibrate and then assess WASABI performance in a controlled and representative setting.

#### Calibration of inference parameters

In the first phase we assessed some critical values to be used in the inference process. We generate realistic GRNs (Fig 2-A) where 20 genes from *in vitro* data were randomly selected with associated *in vitro* estimated parameters (see Supporting information). Interactions were randomly defined in order to create cascade networks with no feedback nor auto-positive feedback as an initial assessment phase.

We limited ourselves to 4 network levels (with 5 genes at each level, see Fig 2-A for an example) because we observed that the information provided by the stimulus is almost completely lost after 4 successive interactions in the absence of positive feedback loops. This is very likely caused by the fact that each gene level adds both some intrinsic noise, due to the bursty nature of gene expression, as well as a filtering attenuation effect due to RNA and protein degradation.

**Fig 2.**
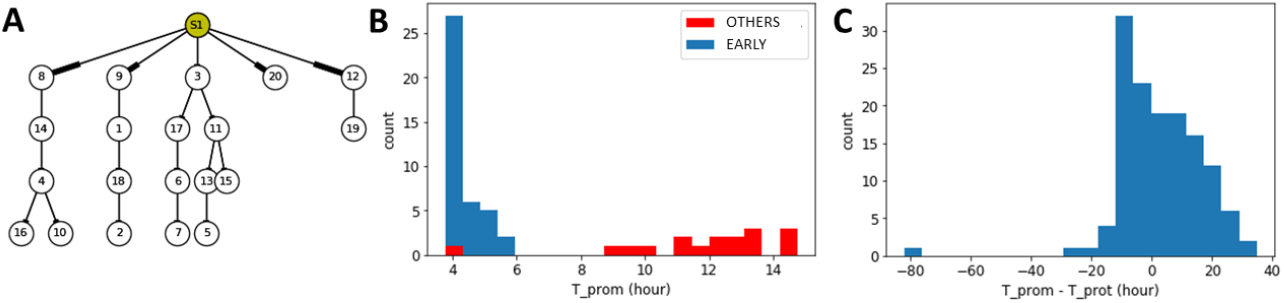
Cascade *in silico* GRN. A) Cascade GRN types are generated to study wave dynamics. Genes correspond to *in vitro* ones with their estimated parameters. S1 corresponds to stimulus. Genes are identified by our list gene ID. B) Based on 10 *in silico* GRN we compare promoter wave time of early genes (blue) with other genes (red). Displayed are promoter waves with a wave time lower than 15h for graph clarity. C) For each interactions of 10 *in silico* GRNs we compute the difference between estimated regulated promoter wave time minus its regulator protein wave time. Distribution of promoter/protein wave time difference is given for all interactions of all *in silico* GRNs.

We first analyzed the special case of early genes that are directly regulated by the stimulus (Fig 2-B). Their promoter wave times were lower than all other genes but one. Therefore we can identify early genes with good confidence, based on comparison of their promoter wave time with a threshold. Given these *in silico* results, we then decided in the WASABI pre-processing step to assume that genes with a promoter wave time below 5h must be early genes, and that genes with a promoter wave time larger than 7h can not be early genes. Interactions between the stimulus and intermediate genes, with promoter wave times between 5h and 7h, have to be tested during the inference iterative process and preserved or not.

We then assessed what would be the acceptable bounds for the difference between regulator protein wave time and regulated gene promoter activity. 10 *in silico* cascade GRNs were generated and simulated for 500 cells to generate population data from which both protein and promoter wave times were estimated for each gene. Based on these data, we computed the difference between estimated regulated promoter wave time minus its regulator protein wave time for all interactions in all networks. The distribution of these wave differences is given in Fig 2-C. One can notice that some wave differences had negative values. This is due to the shape of the Hill interaction function (see eq3 in Method section) with a moderate transition slope (γ = 2). If the protein threshold (which corresponds to typical EC50 value) is too close to the initial protein level, then a slight protein increase will activate target promoter activity. Therefore, promoter activity will be saturated before regulator protein level and thus the difference of associated wave times is negative. This shows that one can accelerate or delay information, depending on the protein threshold value. In order to be conservative during the inference process, we set the RNA/Protein wave difference bounds to [−20h; 30h] in accordance with the distribution in Fig 2-C. One should note that this range, even if conservative, already removes two thirds of all possible interactions, thereby reducing the inference complexity.

We finally observed that for interactions with genes harboring an auto-positive feedback, wave time differences could be larger. In this case, wave difference bounds were estimated to [–30h, 50h] (see supporting information). We interpret this enlargement by an under-sampling time resolution problem since auto-positive feedback results in a sharper transition. As a consequence, promoter state transition from inactive to active is much faster: if it happens between two experimental time points, we cannot detect precisely its wave time.

#### Inference of *in silico* GRNs

WASABI was then tested for its ability to infer *in silico* GRNs (complete definition in supporting information) from which we previously simulated experimental data for mRNA and protein levels at single-cell and population scales. We first assessed the simplest scenario with a toy GRN composed of two branches with no feedback (a cascade GRN; Fig 3-A). The GRN was limited to 6 genes and to 3 levels in order to reduce computational constraints. Nevertheless, even in such a simple case, the inference problem is already a highly complex challenge with more than 10^20^ possible directed networks.

**Fig 3.**
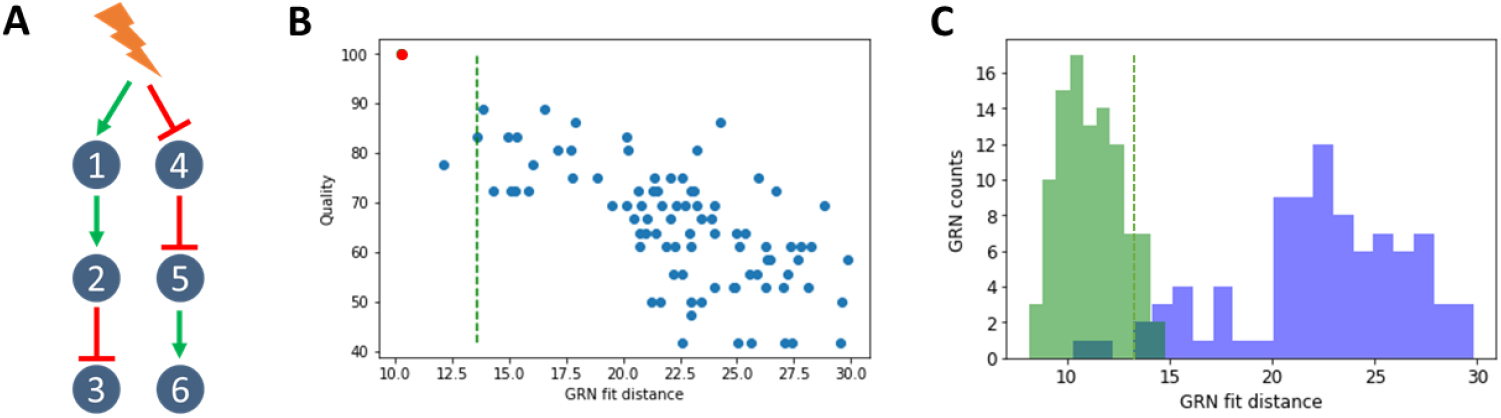
*In silico* cascade GRN inference. A) The cascade GRN. Genes parameters were taken from *in vitro* estimations to mimic realistic behavior. Experimental data were generated to obtain time courses of transciptomic data, at single-cell and population scale, and also proteomic data at population scale. B) WASABI was run to infer *in silico* cascade GRN and generated 88 candidates. A dot represents a network candidate with its associated fit distance and inference quality (percentage of true interactions). True GRN is inferred (red dot, 100% quality). Acceptable maximum fit distance (green dashed line) corresponds to variability of true GRN fit distance. Its computation is detailed in figure C. 3 GRN candidates (including the true one) have a fit distance below threshold. C) Variability of true GRN fit distance (green dashed line in figures B and C) is estimated as the threshold where 95% of true GRN fit distance is below. Fit distance distribution is represented for true GRN (green) and candidates (blue) for cascade *in silico* GRN benchmark. True GRNs are calibrated by WASABI directed inference while candidates are inferred from non-directed inference. Fit distance represents similitude between candidates generated data and reference experimental data.

Wave times were estimated for each gene from simulated population data for RNA and protein (data available in supporting information). Table 1 provides estimated waves time for the cascade GRN. It is clear that the gene network level is correctly reproduced by wave times.

We then ran WASABI on the generated data and obtained 88 GRN candidates (Fig 3-B). The huge reduction in numbers (from 10^20^ to 88) illustrates the power of WASABI to reduce complexity by applying our waves-based constraints. We defined two measures for further assessing the relevance of our candidates:

1. *Quality* quantifies proportion of real interactions that are conserved in the candidate network (see supporting information for a detailed description). A 100% corresponds to the true GRN.

2. A *fit distance*, defined as the mean of the 3 worst gene fit distances, where gene fit distance is the mean of the 3 worst Kantorovitch distances [42] among time points (see the Methods section).

**Table 1.** Wave times. Promoter and protein wave times (in hours) estimated from *in silico* simulated data.

| GRN | Gene | $W_{promoter}$ | $W_{protein}$ |
| --- | --- | --- | --- |
| Cascade | 4 | 4.12 | 12.99 |
|  | 1 | 4.26 | 22.33 |
|  | 5 | 15.19 | 45.50 |
|  | 2 | 17.67 | 44.88 |
|  | 3 | 37.88 | 60.10 |
|  | 6 | 40.06 | 60.72 |

We observed a clear trend that higher quality is associated with a lower fit distance (Fig 3-B), which we denote as a good specificity. When inferring *in vitro* GRNs, one does not have access to quality score, contrary to fit distance. Hence, having a good specificity enables to confidently estimate the quality of GRN candidates from their fit distance. Thus, this result demonstrates that our fit distance criterion can be used for GRN inference. Nevertheless, even in the case of a purely *in silico* approach, quality and fit distance can not be linked by a linear relationship. In other words, the best fit distance can not be taken for the best quality (see below for other toy GRNs). This is likely to be due to both the stochastic gene expression process as well as the estimation procedure. We therefore needed to estimate an acceptable maximum fit distance threshold for true GRN. For this, we ran directed inferences, where WASABI was informed beforehand of the true interactions, but calibration was still run to calibrate interaction parameters. We ran 100 directed inferences and defined the maximum acceptable fit distance (Fig 3-C) as the distance for which 95% of true GRN fit distance was below. This threshold could also be used as a pruning threshold (green dashed line in Fig 3-B) in subsequent iterative inferences, thereby progressively reducing the number of acceptable candidates. We then analyzed a situation where we added either an auto-activation loop or a negative feedback (Fig 4-A and C and supporting information for estimated wave times).

In both cases, GRN inference specificity was lower than for cascade network inference. Nevertheless in both cases the true network was inferred and ranked among the first candidates regarding their fit distance (Fig 4-B and D), demonstrating that WASABI is able to infer auto-positive and negative feedback patterns. However there were more candidates below the acceptable maximum fit distance threshold and there was no obvious correlation between high quality and low fit distance. We think it could be due to data under-sampling regarding the network dynamics (see upper and discussion).

### In vitro application of WASABI

We then applied WASABI on our *in vitro* data, which consists in time stamped single-cell transcriptomic [37] and bulk proteomic data [?] acquired during T2EC differentiation [38], to propose relevant GRN candidates.

We first estimated the wave times (Fig 5). Promoter waves ranged from very early genes regulated before 1h to late genes regulated after 60h. Promoter activity appeared bimodal with an important group of genes regulated before 20h and a second group after 30h. Protein wave distribution was more uniform from 10h to 60h, in accordance with a slower dynamics for proteins. Remarkably, 10 genes harbored non-monotonous evolution of their promoter activity with a transient increase. It can be explained by the presence of a negative feedback loop or an incoherent feed-forward interaction. These results demonstrate that real *in vitro* GRN exhibits distinguishable “waves”.

**Fig 4.**
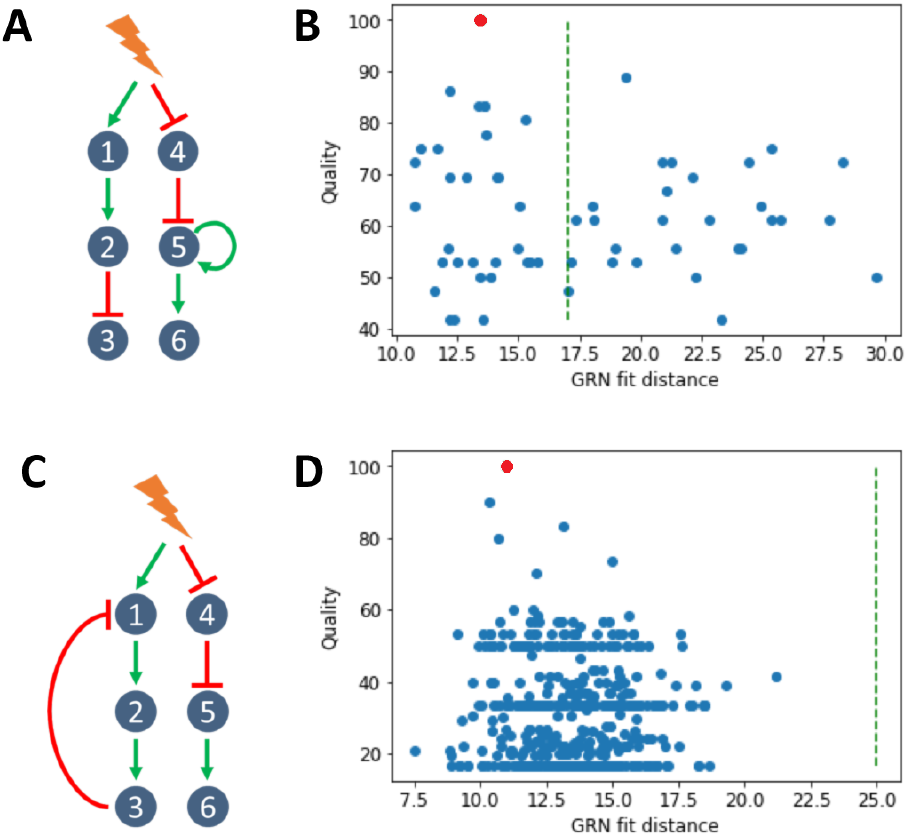
*In silico* GRN with feedbacks. A) Addition of one positive feedback onto the cascade GRN. B) WASABI was run to infer *in silico* cascade GRN with a positive feedback and generated 59 candidates, 31 of which having an acceptable fit distance. See legend to Fig 3-B for details. C) Addition of one negative feedback onto the cascade GRN. D) WASABI was run to infer *in silico* cascade GRN with a negative feedback and generated 476 candidates, all of which having an acceptable fit distance. See legend to Fig 3-B for details.

In order to limit computation time, we decided to further restrict the inference to the most important genes in term of the dynamical behavior of the GRN. We first detected 25 genes that are defined as early with a promoter time lower than 5h. We then defined a second class of genes called “readout” which are influenced by the network state but can not influence in return other genes. Their role for final cell state is certainly crucial, but their influence on the GRN behavior is nevertheless limited. 41 genes were classified as readout so that 24 genes were kept for iterative inference, in addition to the 25 early genes. 9 of these 24 genes have 2 waves due to transient increase, which means that we have 33 waves to iteratively infer.

**Fig 5.**
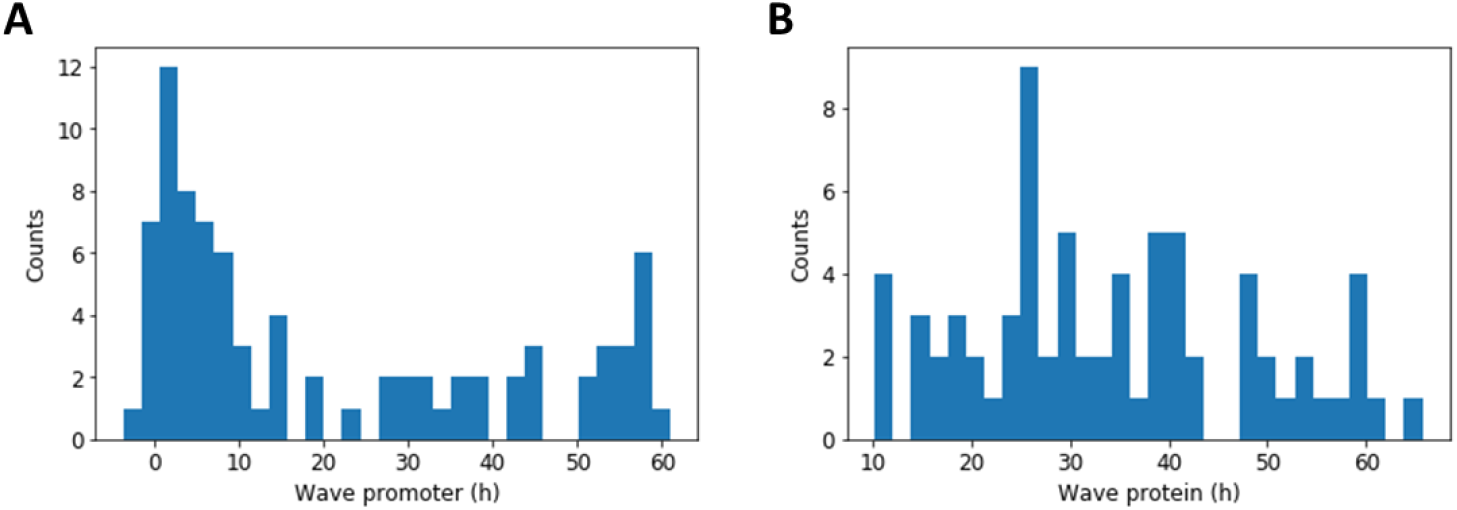
Promoter and protein wave time distributions. Distribution of in vitro promoter (A) and protein (B) wave times for all genes estimated from RNA and proteomic data at population scale. Counts represent number of genes. Note: a gene can have several waves for its promoter or protein.

#### In vitro GRN candidates

After running for 16 days using 400 computational cores, WASABI returned a list of 381 GRN candidates. Candidate fit distances showed a very homogeneous distribution (see supporting information) with a mean value around 30, together with outliers at much higher distances. Removing those outliers left us with 364 candidates. Compared to inference of *in silico* GRN, *in vitro* fitting is less precise, as we could expect. But it is an appreciable performance and it demonstrates that our GRN model is relevant.

We then analyzed the extent of similarities among the GRN candidates regarding their topology by building a consensus interaction matrix (Fig 6-A). The first observation is that the matrix is very sparse (except for early genes in first raw and auto-positive feedbacks in diagonal) meaning that a sparse network is sufficient for reproducing our *in vitro* data. We also clearly see that all candidate GRNs share closely related topologies. This is clearly obvious for early genes and auto-positive feedbacks. Columns with interaction rates lower than 100% correspond to latest integrated genes in the iterative inference process with gene index (from earlier to later) 70, 73, 89, 69 and 29. Results from existing algorithms are usually presented in such a form, where the percent of interactions are plotted [27–29, 35]. But one main advantage of our approach is that it actually proposes real GRN candidates, which may be individually examined.

We therefore took a closer look at the “best” candidate network, with the lowest Fit distance to the data (Fig 6-B). We observed very interesting and somewhat unexpected patterns:

1. Most of the genes (84%) with an auto-activation loop. As mentioned earlier, this was a consensual finding among the candidate networks. It is striking because typical GRN graphs found in the literature do not have such predominance of auto-positive feedbacks.

2. A very large number of genes were found to be early genes that are under the direct control of the stimulus. It is noticeable that most of them were found to be inhibited by the stimulus, and to control not more than one other gene at one next level.

3. We previously described the genes whose product participates in the sterol synthesis pathway, as being enriched for early genes [37]. This was confirmed by our network analysis, with only one sterol-related gene not being an early gene.

4. Among 7 early genes that are positively controlled by the stimulus, 6 are influenced by an incoherent feedforward loop, certainly to reproduce their transient increase experimentally observed [37].

5. One important general rule is that the network depth is limited to 3 genes. One should note that this is not imposed by WASABI which can create networks with unlimited depth. It is consistent with our analysis on signal propagation properties in *in silico* GRN. If network depth is too large, signal is too damped and delayed to accurately reproduce experimental data.

6. One do not see network hubs in the classical sense. The genes in the GRNs are connected to at most four neighbors. The most impacting “node” is the stimulus itself.

7. One can also observe that the more one progress within the network, the less consensual the interaction are. Adding the leaves in the inference process might help to stabilize those late interactions.

Altogether those results show the power of WASABI to offer a brand-new vision of the dynamical control of differentiation.

## Discussion

In the present work we introduced WASABI as a new iterative approach for GRN inference based on single-cell data. We benchmarked it on a representative *in silico* environment before its application on *in vitro* data.

**Fig 6.**
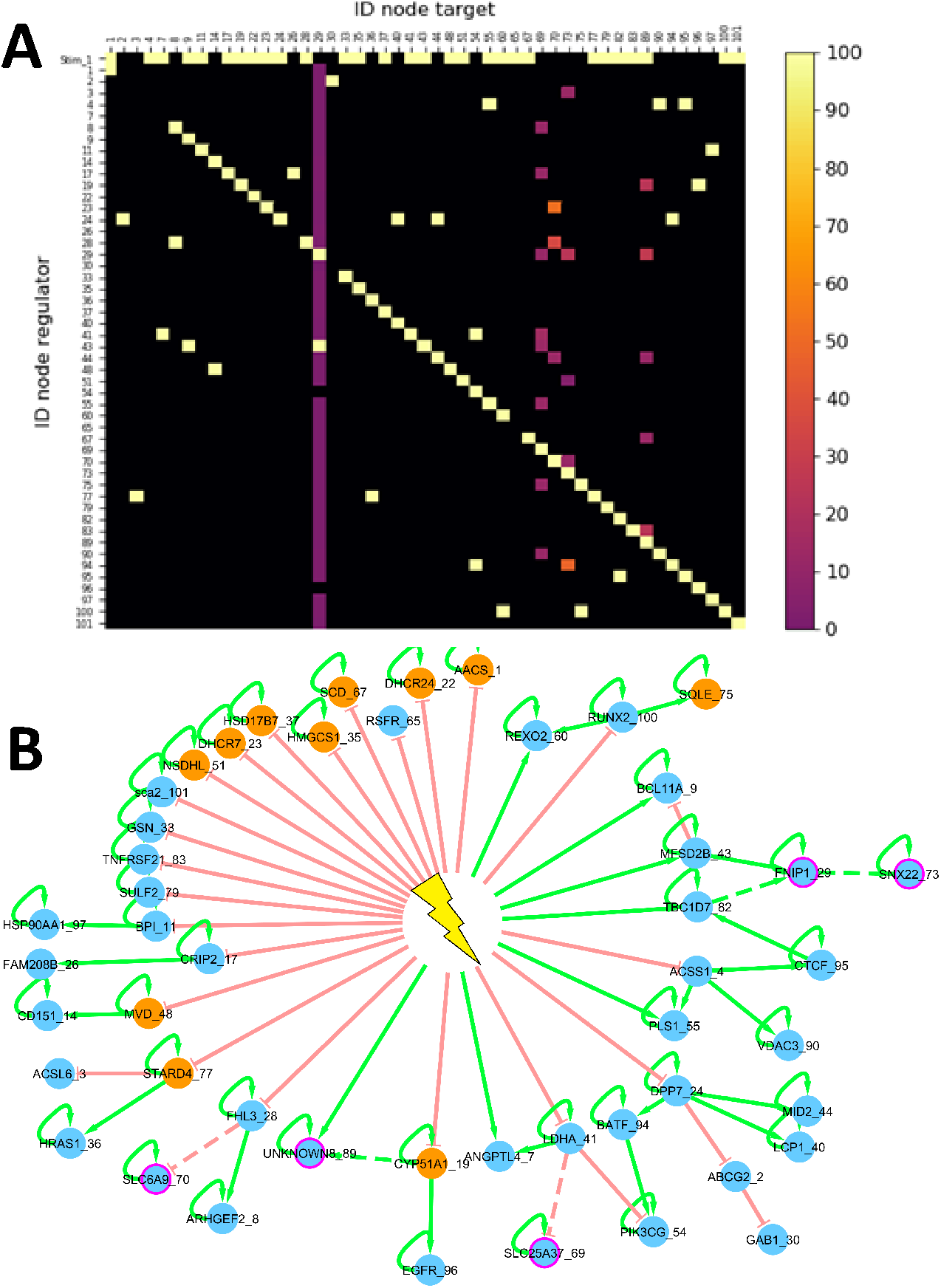
Inference from in vitro data. A) *In vitro* interaction consensus matrix. Each square in the matrix represents either the absence of any interaction, in black, or the presence of an interaction, the frequency of which is color-coded, between the considered regulator ID (row) and regulated gene ID (column). First row correspond to stimulus interactions. B) Best candidate. Green: positive interaction; red: negative interaction; plain lines: interactions found in 100% of the candidates; dashed lines: interaction found only in some of the candidates; orange: genes the product of which participates to the sterol synthesis pathway; purple: 5 last added genes during iterative inference.

### WASABI tackles GRN inference limitations

We are convinced that WASABI has the ability to tackle some general GRN inference issues.

1. WASABI goes beyond mere correlations to infer causalities from time stamped data analysis as demonstrated on *in silico* benchmark (Fig 3) even in the presence of circular causations (Fig 4), based upon the principle that the cause precedes the effect.

2. Contrary to most GRN inference algorithms [27–29, 35] based upon the inference of interactions, WASABI is network centered and generates several candidates with explicitly defined networks topology (Fig 6-B), which is required for prediction making and simulation capability. Generating a list of interactions and their frequency from such candidates is a trivial task (Fig 6-A) whereas the reverse is usually not possible. Moreover, WASABI explicitly integrates the presence of an external stimulus, which surprisingly is never modeled in other approaches based on single-cell data analysis. It could be very instrumental for simulating for example pulses of stimuli.

3. WASABI is not restricted to transcription factors (TFs). Most of the *in vitro* genes we modeled are not TFs. This is possible thanks to the use of our mechanistic model [36] which integrates the notion of timescale separation. It assumes that every biochemical reaction such as metabolic changes, nuclear translocations or post-translational modifications are faster than gene expression dynamics (imposed by mRNA and protein half-life) and that they can be abstracted in the interaction between 2 genes. Our interaction model is therefore an approximation of the underlying biochemical cascade reactions. This should be kept in mind when interpreting an interaction in our GRN: many intermediaries (fast) reactions may be hidden behind this interaction.

4. Optionally, WASABI offers the capability to integrate proteomic data to reproduce translational or post-translational regulation. Our proteomic data [39] demonstrate that nearly half of detected genes exhibit mRNA/protein uncoupling during differentiation and allowed to estimate the time evolution of protein production and degradation rates. Nevertheless, we are not fully explanatory since we do not infer causalities of these parameters evolution. This is a source of improvement discussed later.

5. We deliberately developed WASABI in a “brute force” computational way to guarantee its biological relevance and versatility. This allowed to minimize simplifying assumptions potentially necessary for mathematical formulations. During calibration, we used a simple Euler solver to simulate our networks within model (1). This facilitates addition of any new biological assumption, like post-translation regulations, without modifying the WASABI framework, making it very versatile. Thanks to the splitting and parallelization allowed by WASABI original gene-by-gene iterative inference process, the inference problem becomes linear regarding the network size, whereas typical GRN inference algorithms face combinatorial curse. This strategy also allowed the use of High Parallel Computing (HPC) which is a powerful tool that remains underused for GRN inference [23, 43].

### WASABI performances, improvements and next steps

WASABI has been developed and tested on an *in silico* controlled environment before its application on *in vitro* data. Each *in silico* network true topology was successfully inferred. Cascade type GRN is perfectly inferred (Fig 3) with an excellent specificity. Auto-positive and negative feedback networks (Fig 4) were also inferred, demonstrating WASABI’s ability to infer circular causations, but specificity is lower. This might be due to a time sampling of experimental data being longer than the network dynamic time scale. Auto-positive feedback creates a switch like response, the dynamic of which is much quicker than simple activation. Thus, to capture accurately auto-positive feedback wave time, we should use high frequency time sample for RNA experimental data during auto-positive feedback activation short period. For negative feedback interactions, WASABI calibrated initial increase considering only first experimental time points before feedback effect. Consequently, precision of first interaction was decreased and more false positive sub-GRN candidates were selected. Increasing the frequency of experimental time sampling during initial phase should overcome this problem.

As it stands our mechanistic model is only accounting for transcriptional regulation through proteins. It does not take into account other putative regulation level, including translational or post-translational regulations, or regulation of the mRNA half-life, although there is ample evidence that such regulation might be relevant [44, 45]. Provided that sufficient data is available, it would be straightforward to integrate such information within the WASABI framework. For example, the estimation of the degradation rates at the single-cell level for mRNAs and proteins has recently been described [46], the distribution of which could then be used as an input into the WASABI inference scheme.

Cooperativity and redundancies are not considered in the current WASABI framework, so that a gene can only be regulated by one gene, except for negative feedback or incoherent feedforward interactions. However, many experimentally curated GRN show evidence for cooperations (2 genes are needed to activate a third gene) or redundant interactions (2 genes independently activating a third gene) [47]. We intentionally did not considered such multi-interactions because our current calibration algorithm relies on the comparison of marginal distributions which are not sufficiently informative for inferring cooperative effects. It is our belief that the use of joint distribution of two genes or more should enable such inference. We previously developed in our group a GRN inference algorithm which is based on joint distribution analysis [36] but which does not consider time evolution. We are therefore planning to integrate joint-distribution-based analyses within the WASABI framework in order to improve calibration, by upgrading the objective function with measurement considering joint-distribution comparison.

HPC capacities used during iterative inference impacts WASABI accuracy. Indeed late iterations are supposed more discriminative than the first one because false GRN candidates have accumulated too many wrong interactions so that calibration is not able to compensate for errors. However, if the expansion phase is limited by available computational nodes, the true candidate may be eliminated because at this stage inference is not discriminative enough. Therefore improving computing performances would represent an important refinement and we have initiated preliminary studies in that direction [43].

Nevertheless, despite all possible improvements, GRN inference will remain *per se* an asymptotically solvable problem due to inferability limitations [48], intrinsic biological stochasticity, experimental noise and sampling. This is why we propose a set of GRN candidates with acceptable confidence level. A natural companion of the WASABI approach would be a phase of design of experiments (DOE) specifically aiming at selecting the most informative experiments to discriminate among the candidates. Such DOE procedures have already been developed for GRN inference, but none of them takes into account the mechanistic aspects and the stochasticity of gene expression [48, 49]. Extending the DOE framework to stochastic models is currently being developed in our group.

### New insights on typical GRN topology

The application of WASABI on our *in vitro* model of differentiation generated several GRN candidates with a very interesting consensus topology (Fig 6).

1. We can see that the stimulus (i.e. medium change [37]) is a central regulator of our GRN. We are strongly confident with this result because initial RNA kinetic of early genes can only be explained by fast regulation at promoter level several minutes after stimulation. Proteins dynamics are way too slow to justify these early variations.

2. 22 of the 29 inferred early genes are inhibited by the stimulus, while inhibitions are only present in 7 of the 28 non-early interactions. Thus inhibitions are overrepresented in stimulus-early genes interactions. An interpretation is that most of genes are auto-activated and their inhibition requires a strong and long enough signal to eliminate remaining auto-activated proteins. A constant and strong stimulus should be very efficient for this role like in [32] where stimulus long duration and high amplitude is required to overcome an auto-activation feedback effect. It could be very interesting in that respect to assess how the network would respond to a temporary stimulus, mimicking the commitment experiment described in [37] or [50].

3. None of our GRN candidates do contain so-called “hubs genes” affecting in parallel many genes, whereas existing GRN inferred generally present consequent hubs [26, 28, 29, 35]. A possible interpretation is that hub identifications is mostly a by-product of correlation analysis. This interpretation is in line with the sparse nature of our candidate networks, as compared to some previous network (see e.g. [25] or [51]). This strongly departs with the assumption that small-world network might represent “universal laws” [52].

4. In order to reproduce non-monotonous gene expression variations, WASABI inferred systematically incoherent feedforward pattern instead of “simpler” negative feedback. This result is interesting because nothing in WASABI explain this bias since *in silico* benchmarking proved that WASABI is able to infer simple negative feedbacks (Fig 4). Such “paradoxical components” have been proposed to provide robustness, generate temporal pulses, and provide fold-change detection [53].

5. WASABI candidates are limited in network depth by a maximum of 3 levels. We did not include readout genes during inference but addition of these genes would only increase GRN candidate depth by one level. GRN realistic candidates depth are thus limited by 4 levels. This might be due to the fact that information can only be relayed by limited number of intermediaries because of induced time delay, damping and noise. Indeed, general mechanism of molecules production/degradation behaves exactly as a low pass filter with a cutting frequency equivalent to the molecule degradation rate. Furthermore, protein information will be transmitted at the promoter target level by modulation of burst size and frequency, which are stochastic parameters, thereby adding noise to the original signal.

Such a strong limitation for information carrying capacity in GRN is at stake with long differentiation sequences, say from the hematopoietic stem cell to a fully committed cell. In such a case, tens of genes will have to be sequentially regulated. This might be resolved by the addition of auto-positive feedbacks. Such auto-positive feedbacks will create a dynamic memory whereby the information is maintained even in the absence of the initial information. An important implication is the loss of correlation between auto-activated gene and its regulator gene. Consequently, all algorithms based on stationary RNA single-cell correlation [26, 27] will hardly catch regulators of auto-activated genes.

Considering the importance of auto-positive feedback benefits on GRN information transfert, it is therefore not surprising to see that more than 80% of our GRN genes present auto-positive feedback signatures in their RNA distribution. Moreover, experimentally observed auto-positive feedback influence is stronger in our *in vitro* model than in our *in silico* models. Such a strong prevalence of auto-positive feedbacks has also been observed in a network underlying germ cell differentiation [51]. As mentioned earlier, care should be taken in interpreting such positive influences, which very likely rely on indirect influences, like epigenomic remodeling.

## Materials and Methods

### Mechanistic GRN model

Our approach is based on a mechanistic model that has been previously introduced in [36] and which is summed-up in Fig 7.

**Fig 7.**
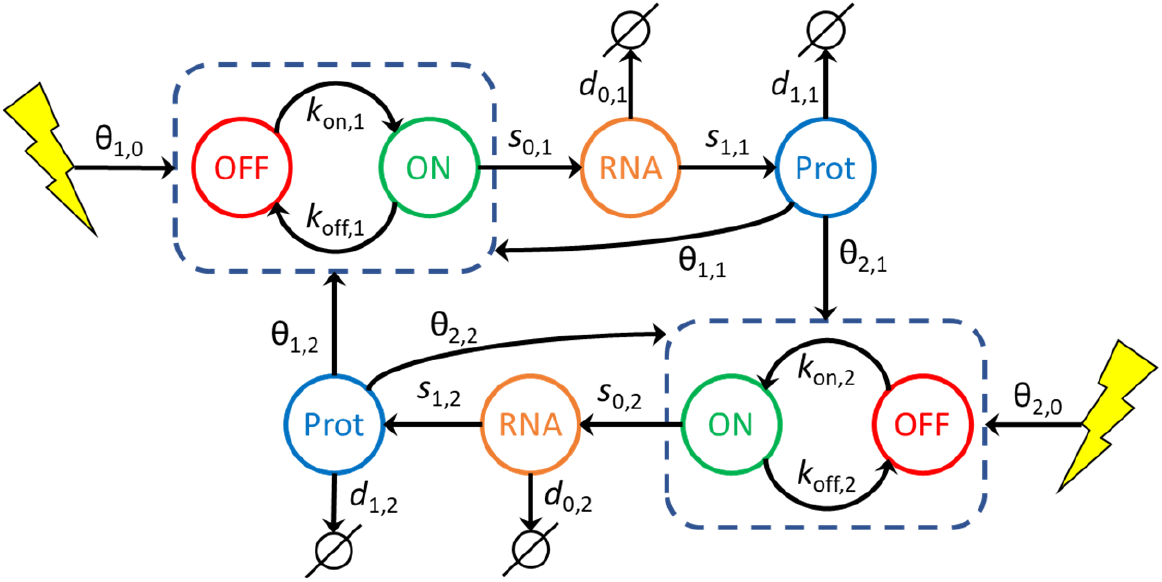
GRN mechanistic and stochastic model. Our GRN model is composed of coupled piecewise deterministic Markov processes. In this example 2 genes are coupled. A gene *i* is represented by its promoter state (dashed box) which can switch randomly from ON to OFF, and OFF to ON, respectively at *k*_on,*i*_ and *k*_off,*i*_ mean rate. When promoter state is ON, mRNA molecules are continuously produced at a *s*_0,*i*_ rate. mRNA molecules are constantly degraded at a *d*_0,*i*_ rate. Proteins are constantly translated from mRNA at a *s*_1,*i*_ rate and degraded at a *d*_1,*i*_ rate. The interaction between a regulator gene *j* and a target gene *i* is defined by the dependence of *k*_on,*i*_ and *k*_off,*i*_ with respect to the protein level *P*_*j*_ of gene *j* and the interaction parameter *θ*_*i*,*j*_. Likewise, a stimulus (yellow flash) can regulate a gene *i* by modulating its *k*_on,*i*_ and *k*_off,*i*_ switching rates with interaction parameter *θ*_*i*,0_.

In all that follows, we consider a set of *G* interacting genes potentially influenced by a stimulus level *Q.* Each gene *i* is described by its promoter state *E_i_ =* 0 (off) or 1 (on), its mRNA level *M*_*i*_ and its protein level *P*_*i*_. We recall the model definition in the following equation, together with notations that will be extensively used throughout this article.

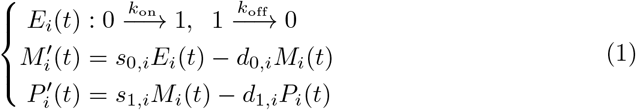

The first line in model (1) represents a discrete, Markov random process, while the two others are ordinary differential equations (ODEs) describing the evolution of mRNA and protein levels. Interactions between genes and stimulus are then characterized by the assumption that *k*_on_ and *k*_off_ are functions of *P* = (*P*_1_,…, *P*_*G*_) and *Q.* The form for *k*_*on*_ is the following (for *k*_off_, replace *θ*_*i,j*_ by –*θ*_*i,j*_):

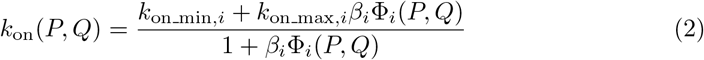

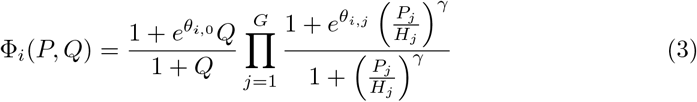

This interaction function slightly differs from [36] since auto-feedback is considered as any other interactions and stimulus effect is explicitly defined. Exponent parameter γ is set to default value 2. Interaction threshold *H*_*j*_ is associated to protein *j.* Interaction parameters *θ*_*i,j*_ will be estimated during the iterative inference. Parameter *β*_*i*_ corresponds to GRN external and constant influence on gene to define its basal expression: it is computed at simulation initialization in order to set *k*_on_ and *k*_off_ to their initial value. From now on, we drop the index *i* to simplify our notation when there is no ambiguity.

### Overview of WASABI workflow

WASABI framework is divided in 3 main steps. First, individual gene parameters defined in model (1) (all except *θ* and *H*) are estimated before network inference from a number of experimental data types acquired during T2EC differentiation. They include time stamped single-cell transcriptomic [37], bulk transcription inhibition kinetic [37] and bulk proteomic data [39]. In a second step, genes are sorted regarding their wave times (see “Results” section for a description of wave concept) estimated from the mean of single cell transcriptomic data for promoter waves, and bulk proteomic data for protein waves. Finally, network iterative inference step is performed from single transcriptomic data, previously inferred gene parameters and sorted genes list. All methods are detailed in following sections, an overview of workflow is given by Fig 8.

For T2EC *in vitro* application, tables of gene parameters and wave times are provided in supporting information. For *in silico* benchmarking we assume that gene parameters *d*_0_, *d*_1_, *s*_1_ are known. Single-cell data and bulk proteomic data are simulated from *in silico* GRNs for time points 0, 2, 4, 8, 24, 33, 48, 72 and 100h.

**Fig 8.**
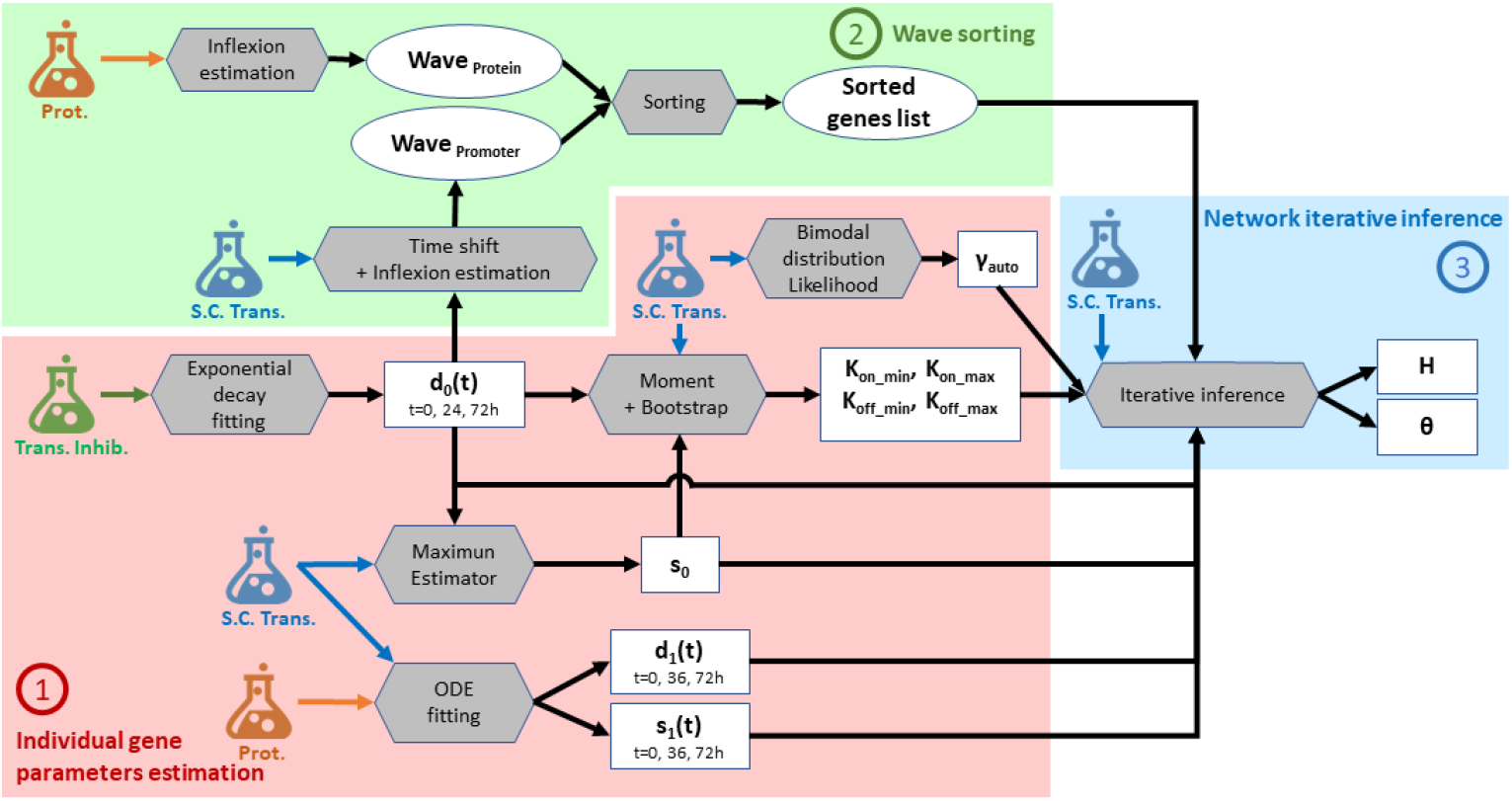
Parameters estimation workflow. Schematic view of WASABI workflow with 3 main steps: (1) individual gene parameters estimation (red zone), (2) waves sorting (green zone) and (3) network iterative interaction inference (blue zone). Wave concept is introduced in “Result” section. Model parameters (square boxes) are estimated from experimental data (flasks) with a specific method (grey hexagones). All methods are detailed in “Method” section. Estimated data relative to waves are represented by round boxes. Input arrows represent data required by methods to compute parameters. There are 3 types of experimental data, (i) bulk transcription inhibition kinetic (green flask), (ii) single-cell transcriptomic (blue flask) and (iii) proteomic data (orange flask). Model parameters are specific to each gene, except for *θ*, which is specific to a pair of regulator/regulated genes. Notations are consistent with Eq(1), *γ*_*auto*_ represents exponent term of auto-positive feedback interaction. Only *d*_0_(*t*), *d*_1_(*t*) and *s*_1_(*t*) are time dependent. One gene can have several wave times.

### First step - Individual gene parameters estimation

#### Exponential decay fitting for mRNA degradation rate (*d*_0_) estimation

The degradation rate *d*_0_ corresponds to active decay (i.e. destruction of mRNA) plus dilution due to cell division. The RNA decay was already estimated in [37] before differentiation (0h), 24h and 72h after differentiation induction from population-based data of mRNA decay kinetic using actinomycin D-treated T2EC (osf.io/k2q5b). Cell division dilution rate is assumed to be constant during the differentiation process and cell cycle time has been experimentally measured at 20h [38].

#### Maximum estimator for mRNA transcription rate (s_0_) estimation

To infer the transcription rate *s*_0_, we used a maximum estimator based on single-cell expression data generated in [37]. We suppose that the highest possible mRNA level is given by *s*_0_*/d*_0_. Thus *s*_0_ corresponds to the maximum mRNA count observed in all cells and time points multiplied by 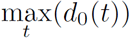.

#### Method of moments and bootstrapping for range of promoter switching rates (*k*_on/off_min/max_) estimation

Dynamic parameters *k*_on_ and *k*_off_ are bounded respectively by constant parameters [*k*_on_min_;*k*_on_max_] and [*k*_off_min_; *k*_off_max_] (see Eq (2)) which are estimated as follows from time course single-cell transcriptomic data. Parameters *s*_0_ and *d*_0_(*t*) are supposed to be previously estimated for each gene at time *t.*

Range parameters shall be compliant with constraints (Eq (4)) imposed by the transcription dynamic regime observed *in vitro.* RNA distributions [37] have many zeros, which is consistent with the bursty regime of transcription. There is no observed RNA saturation in distributions. Moreover, all GRN parameters should also comply with computational constraints. On the one hand, the time step *dt* used for simulations shall be small enough regarding GRN dynamics to avoid aliasing (under-sampling) effects. On the other hand, *dt* should not be too small to save computation time. These constraints correspond to

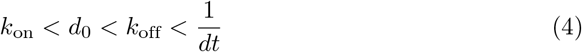

and we deduce inequalities for ranges:

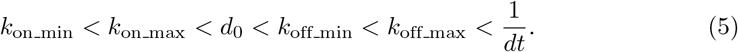

We set the default value *k*_on___min_ to 0.001 h^−1^. Parameter *k*_on___max_ is estimated from time course single-cell transcriptomic data after removing zeros. This truncation mimics a distribution where gene is always activated, so that *k*_on_ is close to its maximum value *k*_on_max_. With these truncated distributions, for each time point *t*, we estimate *k*_on,*t*_ using a moment-based method defined in [54]. We bootstrapped 1000 times to get a list of *k*_on,*t,n*_ with index *n* corresponding to bootstrap sample *n.* For each time point we compute the 95% percentile of *k*_on,*t,n*_, then we consider the mean value of these percentiles to have a first estimate of *k*_on_max_. This *k*_on_max_ is then down and up limited respectively between *k*_on___max_lim_min_ and *k*_on_max_lim_max_ given in Eq (6) to guarantee that observed *k*_on_ can be easily reached during simulations with reasonable values of protein level (because of asymptotic behavior of interaction function). In other words *k*_on_max_ shall not be too close from minimum or maximum observed *k*_on_ considering 10% margins. Finally, this limited *k*_on_max_ is up-limited by 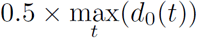 to guarantee a 50% margin with *d*_0_(*t*).

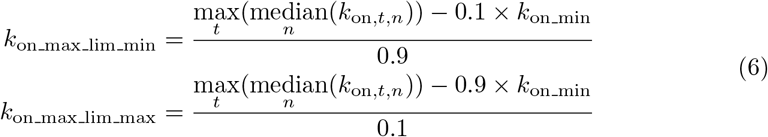

Parameter *k*_off_min_ is set to 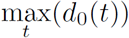 to comply with equation Eq (5). Parameter *k*_off_max_ is estimated like *k*_on_max_ from time course single-cell transcriptomic data but without zero truncation. For each time point *t*, we estimate *k*_off,*t*_ using a moment-based method defined in [54]. We bootstrapped 1000 times to get a list of *k*_off,*t,n*_ with index *n* corresponding to bootstrap sample n. For each time point we compute the 95% percentile of *k*_off,*t,n*_, then we consider the mean value of these percentiles to have a first estimate of *k*_off max_. This *k*_off_max_ is then down and up limited respectively between *k*_off_max_lim_min_ and *k*_off_max_lim_max_ given in Eq (7) to guarantee that observed *k*_off_ can be easily reached during simulations with reasonable values of protein level (because of asymptotic behavior of interaction function). In other words *k*_off_max_ shall not be too close from minimum or maximum observed *k*_off_ considering 10% margins. Finally, this limited *k*_off_max_ is up-limited by 1/*dt* to guaranty simulation anti-aliasing.

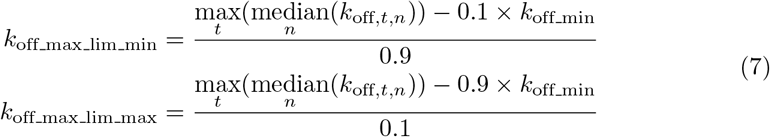

#### ODE fitting for protein translation and degradation rates (*d*_1_,*s*_1_) estimation

Rates *d*_1_(*t*) and *s*_1_(*t*) are estimated from comparison of proteomic population kinetic data [39] with RNA mean value kinetic data computed from single-cell data [37]. Parameter *d*_1_(*t*) corresponds to protein active decay rate while total protein degradation rate *d*_1_*tot*_(*t*) includes decay plus cell division dilution. Associated total protein half-life is referred to as *t*_1_*tot*_(*t*). Parameters *s*_1_(*t*) and *d*_1_*tot*_(*t*) are estimated using a calibration algorithm based on a maximum likelihood estimator (MLE) from package [55]. Objective function is given by the Root Mean Squared Error function (provided by the package) comparing experimental protein counts with simulated ones given by ODEs from our model (1) with RNA level provided by experimental mean RNA data:

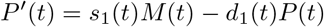

52 out of our 90 selected genes were detected in proteomic data. 23 of these fit correctly experimental data with a constant *d*_1_ and *s*_1_ during differentiation. 5 genes were estimated with a variable *s*_1_(*t*) and a constant *d*_1_ to fit a constant protein level with a decreasing RNA level. For the remaining 24 genes, protein level decreased while RNA is constant, which is modeled with *s*_1_ constant and *d*_1_(*t*) variable.

For the genes that were not detected in our proteomic data we turned to the literature [56] and found 13 homologous genes with associated estimation of *d*_1_ and *s*_1_. For the remaining 25 genes, we estimated parameters with the following rationale: we consider that the non-detection in the proteomic data is due to low protein copy number, lower than 100. Moreover [56] proposed an exponential relation between *s*_1_ and the mean protein level that we confirmed with our data (see supporting information), resulting in the following definition:

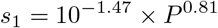

Linear regression was performed using the Python `scipy.stats.linregress()` method from Scipy package with the following parameters: *r*^2^ = 0.55, slope = 0.81, intercept = –1.47 and *p =* 2.97 × 10^−9^. Therefore, if we extrapolate this relation for low protein copy numbers assuming *P* < 100 copies, *s*_1_ should be lower than 1 molecule/RNA/hour. Assuming the relation

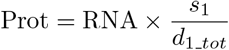

between mean protein and RNA levels, we deduced a minimum value of *d*_1_ from mean RNA level given by: *d*_1_ > RNA/100. We set *s*_1_ and *d*_1_ respectively to their maximum and minimum estimated values.

#### Bimodal distribution likelihood for auto-positive feedback exponent (*γ*_*auto*_) estimation

We inferred the presence of auto-positive feedback by fitting an individual model for each gene, based on [36]. The model is characterized by a Hill-type power coefficient. The value of this coefficient was inferred by maximizing the model likelihood, available in explicit form. The key idea is that genes with auto-positive feedback typically show, once viewed on an appropriate scale, a strongly bimodal distribution during their transitory regime. The interested reader may find some details in the supplementary information file of [36], especially in sections 3.6 and 5.2. Note that such auto-positive feedback may reflect either a direct auto-activation, or a strong but indirect positive loop, potentially involving other genes. Estimated Hill-type power coefficients for *in silico* and *in vitro* networks are provided in supporting information.

### Second step - Waves sorting

#### Inflexion estimator for wave time estimation

Wave time for gene promoter *W*_*prom*_ and protein *W*_*prot*_ are estimated regarding their respective mean trace 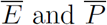. Estimation differs depending on mean trace monotony. *In vitro* wave times are provided in supporting information.

1) If the mean trace is monotonous (checked manually), it is smoothed by a 3rd order polynomial approximation using method `polyld()` from python numpy package. Wave time is then defined as the inflection time point of polynomial function where 50% of evolution between minimum and maximum is reached.

2) If the mean trace is not monotonous, it is approximated by a piecewise-linear function with 3 breakpoints that minimizes the least square error. Linear interpolations are performed using the `polynomial.polyfit()` function from python `numpy` package. Selection of breakpoints is performed using `optimize.brute()` function from python `numpy` package.

We obtained a series of 4 segments with associated breakpoints coordinate and slope. Slopes are thresholded: if absolute value is lower than 0.2 it is considered null. Then, we looked for inflection break times where segments with non null slope have an opposite sign compare to the previous segment, or if previous segment has a null slope. Each inflection break time corresponds to an initial effect of a wave. A valid time, when wave effect applies, is associated and corresponds to next inflection break time or to the end of differentiation. Thus, we obtained couples of inflection break time and valid time which defined the temporal window of associated wave effect. For each wave window, if mean trace variation between inflection break time and valid time is large enough (i.e., greater than 20% of maximal variation during all differentiation process for the gene), a wave time is defined as the time where half of mean trace variation is reached during wave time window.

Protein mean trace 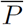 is given by proteomic data if available, else it is computed from simulation traces with 500 cells using the model with the parameters estimated earlier. Promoter mean trace 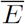 is computed as follows from mean RNA trace (from single-cell transcriptomic data) with time delay correction induced by mRNA degradation rate *d*_0_.

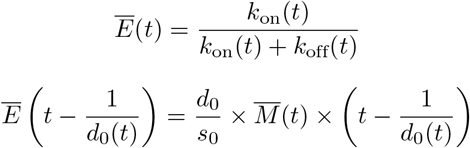

#### Genes sorting

Genes are sorted regarding their promoter waves time *W*_*prom*_. Genes with multiple waves, in case of feedback for example, are present several times in the list. Moreover, genes are classified by groups regarding their position in the network. Genes directly regulated by the stimulus are called the early genes; Genes that regulates other genes are defined as regulatory genes; Genes that do not influence other genes are identified as readout genes. Note that genes can belong to several group.

We can deduce the group type for each gene from its wave time estimation. Subsequent constraints have been defined from *in silico* benchmarking (see Results section). A gene *i* belongs to one of these groups according to following rules:

- if *W*_*prom*_ < 5h then it is an early gene
- if *W*_*prom*_ < 7h then it could be an early gene or another types
- if 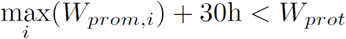 then it is a readout gene
- else it could be a regulatory or a readout gene

### Third step - Network iterative inference

#### Interaction threshold (*H*)

Interaction threshold *H* is estimated for each protein. It corresponds to mean protein level at 25% between minimum and maximum mean protein level observed during differentiation by *in silico* simulations:

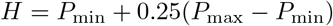

We choose the value of 25% to maximize the amplitude variation of *k*_on_ and *k*_off_ of gene target induced by the shift of the regulator protein level from its minimal to maximal value (see Eq(2)).

#### Iterative calibration algorithm (*θ*_*i,j*_)

The following algorithm gives a global overview of the iterative inference process:

##### Generate_EARLY_network()

In a first step we calibrate the interactions between early genes and stimulus (*θ*_*i*, 0_) to obtain an initial sub-GRN. Calibration algorithm *Calibrate()* is defined below.

##### List_genes_sorted_by_Wave_time

This list is computed prior to iterative inference (see previous subsection).

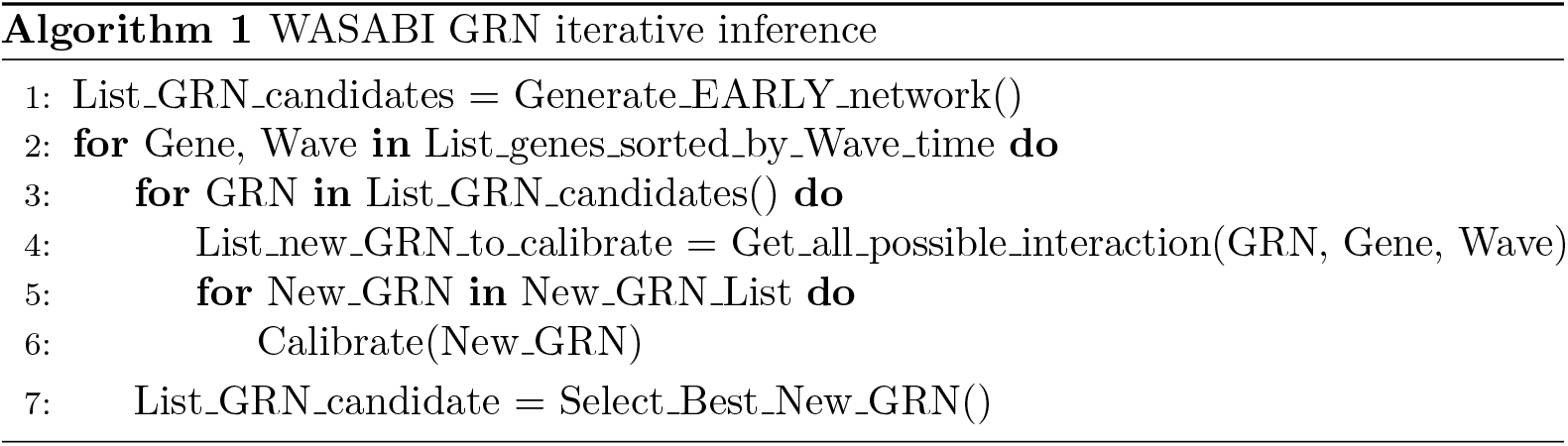

##### Get_all_possible_interaction(GRN, Gene, Wave)

For each GRN candidate we estimate all possible interactions with the new gene and prior regulatory genes, or stimulus, regarding their respective promoter wave and protein wave with the following logic: if promoter wave is lower than 7h, interaction is possible between stimulus and the new gene. If the difference of promoter wave minus protein wave is between −20h and +30h, then there is a possible interaction between the new gene and regulatory gene. Note: if WASABI is run in “directed” mode, only the true interaction is returned.

##### Calibrate(New_GRN)

For interaction parameter calibration we used a Maximum Likelihood Estimator (MLE) from package `spotpy` [55]. The goal is to fit simulated single-cell gene marginal distribution with *in vitro* ones tuning efficiency interaction parameter *θ*_*i*,*j*_. For *in silico* study we defined **GRN Fit distance** as the mean of the 3 worst gene-wise fit distances. For *in vitro* study we defined **GRN Fit distance** as the mean of the fit distances of all genes. Gene-wise fit distance is defined as the mean of the 3 higher Kantorovitch distances [42] among time points. For a given time point and a given gene, the Kantorovitch fit distance corresponds to a distance between marginal distributions of simulated and experimental expression data. At the end of calibration the set of interaction parameter *θ*_*i,j*_ with associated GRN Fit distance is returned.

##### Select_Best_New_GRN()

We fetch all GRN calibration fitting outputs from remote servers and select best new GRNs to be expanded for next iteration updating list of List_GRN_candidate. New networks candidates are limited by number of available computational cores.

### GRN simulation

We use a basic Euler solver with fixed time step (*dt* = 0.5h) to solve mRNA and protein ODEs [36]. The promoter state evolution between *t* and *t + dt* is given by a Bernoulli distributed random variable

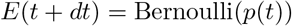

drawn with probability *p(t)* depending on current *k*_on_, *k*_off_ and promoter state:

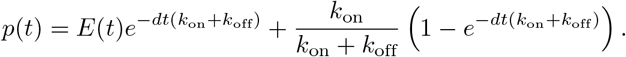

Time-dependent parameters like *d*_0_, *d*_1_ and *s*_1_ are linearly interpolated between 2 points. The stimulus *Q* is represented by a step function between 0 and 1000 at *t* = 0h. Simulation starts at *t* = –60h to ensure convergence to steady state before the stimulus is applied. Parameters *k*_on_ and *k*_off_ are given by Eq (2).

## Author summary

All cells have to make everyday decisions regarding their behavior in response to changing environment. Such decisions result from the dynamical behavior of an underlying gene regulatory network. Inferring the structure of such networks is an inverse problem which has occupied the systems biology community for decades. We propose in the present work a divide-and-conquer strategy called WASABI, which splits the potentially untractable global problem into much simpler subproblems. We show that by adding one gene at a time, we can infer small networks, the behavior of which has been simulated *in silico* using a mechanistic model which incorporates the fundamentally probabilistic nature of the gene expression process. When applied to real-life data, our algorithm sheds a new fascinating light onto the molecular control of a differentiation process. It is our hope that WASABI will prove useful in helping biologists to fully exploit the power of time-stamped single-cell data.

## Acknowledgments

We thank the computational center of IN2P3 (Villeurbanne/France), specially Pascal Calvat, for access to HPC facilities; Eddy Caron (Avalon, ENS Lyon/INRIA) for his support on parallel computing implementation; Patrick Mayeux for proteomic data; and Rudiyanto Gunawan (ETH, Zürich) for critical reading of the manuscript. We would like to thank all members of the SBDM team, Dracula team, and Camilo La Rota (Cosmotech) for enlightening discussions, We also thank the BioSyL Federation and the Ecofect Labex (ANR-11-LABX-0048) of the University of Lyon for inspiring scientific events.

